# Protein tyrosine phosphatase receptor type kappa (PTPRκ) regulates Superior ON-Direction Selective Ganglion Cell development, facilitating image stabilization

**DOI:** 10.64898/2026.01.07.698158

**Authors:** Tzu-Huai Lin, Annika Balraj, Timour Al-Khindi, James Kiraly, Felice A. Dunn, Alex L. Kolodkin

## Abstract

In vertebrates, gaze stabilization during global visual motion requires ON direction-selective ganglion cells (oDSGCs) in the Accessory Optic System (AOS) to drive the optokinetic reflex (OKR). Three types of oDSGCs form independent mosaics; however, the mechanisms that specify and maintain their numbers to create these mosaics during development remain unknown. Here, we identify protein tyrosine phosphatase receptor type kappa (*Ptprk*) as a key regulator of Superior oDSGC density, the subtype tuned to upward motion. High-depth single-cell RNA sequencing (scRNAseq) reveals that *Ptprk* is selectively enriched in Superior oDSGCs compared to Inferior oDSGCs, which detect downward motion. Genetic deletion of *Ptprk* selectively halves the number of Superior oDSGCs while preserving the number of Inferior oDSGCs. Anatomically, Superior oDSGCs in *Ptprk* mutants exhibit compensatory neurite remodeling to preserve mosaics. Physiologically, oDSGCs have broader tuning curves, and a subset of Superior oDSGCS have aberrant preferred directions in *Ptprk* mutants. Postsynaptically, loss of *Ptprk* reduces oDSGC connectivity to central targets. Behaviorally, the OKR responses to upward motion are attenuated in both global and pan-retinal *Ptprk* mutants, confirming a retinal role for *Ptprk* in upward image stabilization. Together, these findings demonstrate that the density of a retinal ganglion cell type is under molecular control, and that reducing density drives neurite remodeling, alters circuit computation, and ultimately impairs visually-guided behavior.

## Introduction

Visual processing begins in the retina, where photons are absorbed by millions of photoreceptors, generating electrical and chemical signals that are then processed by parallel retinal circuits. These circuits extract specific visual features such as contrast, orientation, and motion direction. This information is then sent to central targets via retinal ganglion cells (RGCs), a diverse neuronal population comprising less than one percent of total retinal neurons. Transcriptomic profiling has identified over 45 distinct types of RGCs in mice^1,2^ and at least 23 in humans^3^. The majority of the RGC types tile the retina in a regular array, each forming a mosaic that also serves to identify them as a unique cell type^4,5^. It remains poorly understood how these RGC types are specified and, importantly, maintained with respect to number, spacing, and dendrite morphology such that each can be correctly integrated into their corresponding retinal circuits to evenly sample visual space.

ON-Direction Selective Ganglion Cells (oDGSCs), an integral component of direction-selective (DS) circuits within the Accessory Optic System (AOS), sense slow, directionally biased object motion via greater spike responses to motion in a preferred direction and fewer spikes in the opposite, or null, direction. oDSGCs facilitate image stabilization on the retina during self motion. By encoding and relaying such information for visuovestibular computation, these DS circuits generate the optokinetic reflex (OKR), a reflexive oculomotor response. The OKR consists of slow eye tracking following the direction of motion, interspersed with rapid compensatory saccades in the opposite direction^6,7^. In the retina, oDSGC somas reside in the ganglion cell layer (GCL) and their dendrites stratify in the inner plexiform layer (IPL). Type 5 and 7 bipolar cells (BCs) in the inner nuclear layer (INL) provide excitatory input onto oDSGCs, and amacrine cells (ACs), including starburst amacrine cells (SACs) residing in the INL and GCL, provide inhibitory input to oDSGCs^8^. Direction selectivity depends critically on asymmetric inhibitory input from SACs onto oDSGCs; this inhibition is stronger on the null side and weaker on the preferred side of the oDSGC dendritic arbor, enabling robust firing only when stimuli move in the preferred direction^9,10^. Three major oDSGCs types are each tuned to a specific direction in visual space: Superior (up), Inferior (down) or Nasal (forward) ^8,11,12^. Additional evidence supports the existence of a rare fourth Temporal subtype that senses backward motion^12^. These functional distinctions are reflected in their central projections; Superior oDSGCs project their axons to the dorsal medial terminal nucleus (dMTN), Inferior oDSGCs to ventral MTN (vMTN), and Nasal oDSGCs to both the nucleus of optic tract (NOT) and the dorsal terminal nucleus (DTN) ^11,13,14^.

Despite functional and anatomical distinctions among oDSGCs, previous transcriptomic profiling shows that these oDSGCs are closely related at the level of gene expression. Nevertheless, our high-depth single-cell RNA sequencing (scRNAseq) identified unique transcriptomic profiles that distinguish Superior and Inferior oDSGC populations. This analysis revealed selective expression of the transcription factor *Tbx5* in Superior oDSGCs; loss of *Tbx5* prevents differentiation of the Superior oDSGC population, leading to the complete loss of the vertical OKR response^15^. Our scRNAseq analysis also uncovered additional differentially expressed genes in vertically tuned oDSGCs: Fibrinogen C domain containing 1 (*Fibcd1*) in Inferior oDSGCs, and Protein tyrosine phosphatase receptor kappa (*Ptprk*) in Superior oDSGCs^15^. Both of these genes encode cell surface molecules.

Here, we investigate the role of *Ptprk* in Superior oDSGCs and its contribution to the OKR. We examine *Ptprk* expression oDSGCs and investigate the effect of PTPRκ on Superior oDSGC number, mosaic spacing, and dendritic morphology. We also assess functional parameters, including spike responses and directional tuning in *Ptprk* mutant oDSGCs and evaluate the OKR performance in *Ptprk* mutants. Our results show that *Ptprk* is critical for normal Superior oDSGC numbers, morphology, and for normal image stabilization.

## Results

### *Ptprk* is differentially expressed in Superior oDSGCs

To investigate oDSGC directional tuning, we identified differentially expressed genes in Superior vs. Inferior murine oDSGCs^15^. *Ptprk* is selectively expressed in Superior oDSGCs, and this can be observed between postnatal day (P) 5 to P12, spanning the time period when direction selectivity is established (P6-P8) (Fig. 1A, B). We examined *Ptprk* expression patterns using RNAscope *in situ* hybridization, using *Spig1-GFP*^+^ reporter as a marker for this subclass of oDSGC in the nasal and ventral domains of the retinas^13^, and observed robust *Ptprk* expression in Superior oDSGCs. *Ptprk* was not expressed in Inferior oDSGCs, which we identified using *Fibcd1*, a gene that is expressed in Inferior oDSGCs but not Superior oDSGCs in our scRNAseq data set (Fig. 1C-D). This finding is corroborated by two additional neonatal murine scRNAseq datasets (Rheaume *et al.* for P5 RGCs^16^ and Somaiya *et al.* for P8 RGCs^17^). In the P5 dataset^16^, oDSGCs reside in cluster 32 (C32), characterized by marker genes such as *Pappa*, *Robo1* and *Gpr88*^16,18,19^; parsing RGCs in C32 into Superior, Inferior and Nasal oDSGC subclusters (see Materials and Methods) revealed coexpression of *Ptprk* with *Tbx5*, an essential Superior oDSGC marker^15^, but not with the Inferior oDSGC marker *Fibcd1* (Fig. 1E). A previously generated P56 RGC dataset^1^ also showed *Ptprk* expression in the putative vertical oDSGC cluster (cluster10_Novel), marked by *Nr2f2*^20^, *Bnc2*^21^ and *Syt6*^18^ (Fig. S1A). Further re-clustering of RGCs in cluster 10 from the P56 dataset was hindered by insufficient depth of these sequencing data. Though *Ptprk* is expressed in several types of amacrine cells, critical components of AOS DS circuitry such as starburst amacrine cells (SACs) (*ChAT^+^*) and *vGlut3*^+^ amacrine cells^22^ do not express *Ptprk* (Fig. S1B, C). Furthermore, analysis of a human fetal RGC dataset^3^ reveals that *Ptprk* is expressed in cluster 21, which exhibits a marker profile (*Bnc2^+^*, *Nr2f2^+^*, *Fstl4^+^*, *Robo1^+^* and *Tbx5^+^*)^19^ consistent with that of putative human Superior oDSGCs (Fig. 1F). Taken together, these data suggest that PTPRk expression, and possibly function, in Superior oDSGCs is evolutionarily conserved.

**Figure 1.**
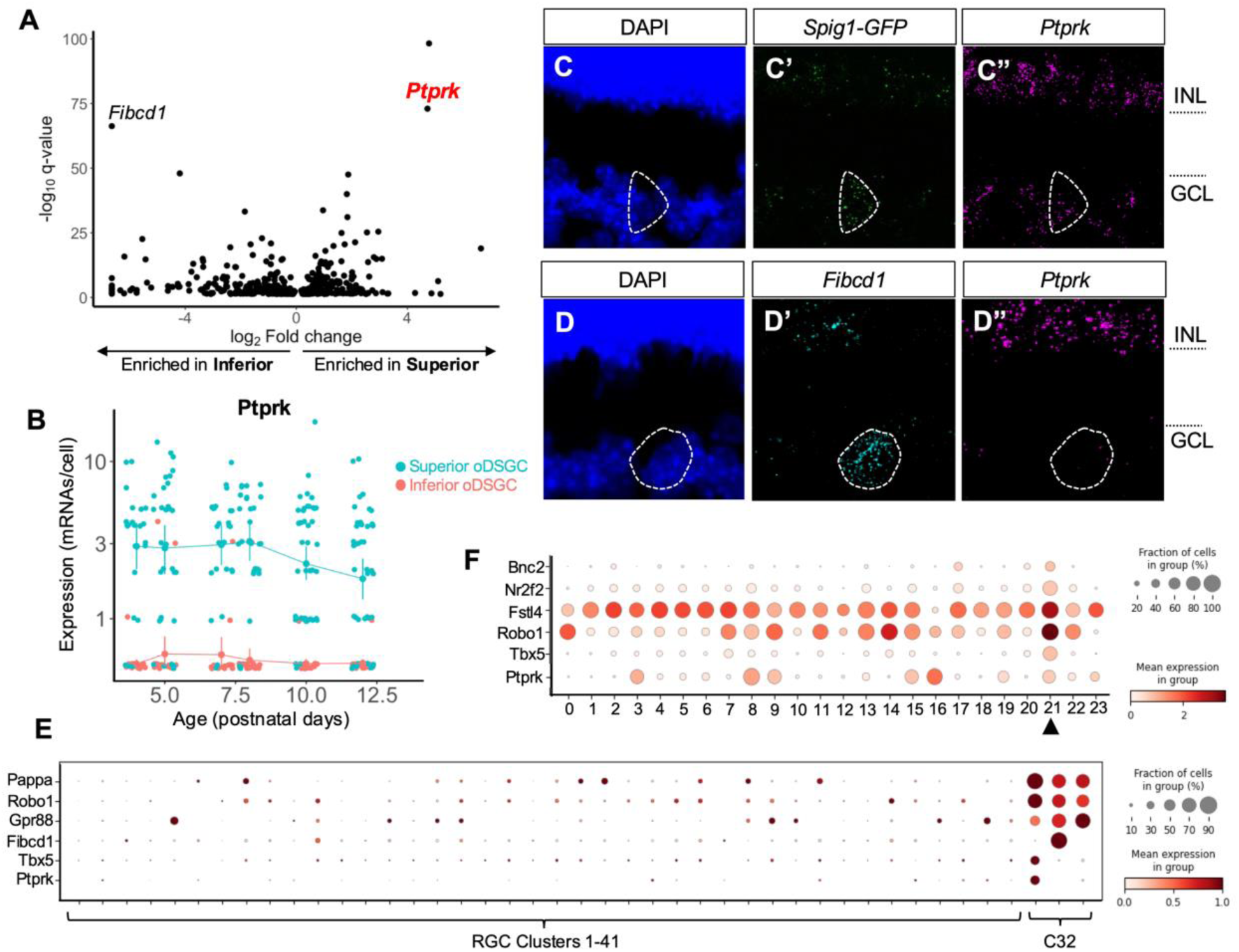
Expression pattern of *Ptprk*. **(A)** A volcano plot generated from the previous work^15^ showing q values and fold change for genes that are significantly differentially expressed between Superior and Inferior oDSGCs. All dots show genes that have q < 0.05. *Ptprk* is differentially expressed in Superior oDSGCs whereas *Fibcd1* is specifically enriched in Inferior oDSGCs. **(B)** *Ptprk* expression (mRNAs/cell) remains higher in Superior oDSGCs from P4 to P12 compared to that in Inferior oDSGCs. Data are presented as mean ± 95% confidence intervals. **(C-C’’, D-D’’)** RNAscope *in situ* hybridization in P8 *Spig1-GFP* retinas shows *Ptprk* expression present in GFP^+^ Superior oDSGCs (C-C’’) but absent in Fibcd1^+^ Inferior oDSGCs (D-D’’). **(E)** Dotplot showing *Ptprk* expression in cluster 32 (C32) of the P5 RGC dataset along with the expression of of oDSGC markers *Pappa*, *Robo1* and *Gpr88*^16,18,19^, Superior oDSGC marker *Tbx5* and Inferior oDSGC marker *Fibcd1*^15^. Reclustering of C32 of the P5 dataset is described in our previous work^19^. **(F)** Dotplot of oDSGC markers shows the specificity of *Ptprk* expression in cluster 21, which contains putative oDSGCs. The analysis is described in the previous work^19^.

### *Ptprk* is required for maintaining the density, but not regular spacing, of Superior oDSGCs mosaics

To investigate the role of *Ptprk* in Superior oDSGCs development, we used *Spig1-GFP* to quantify the number of Superior oDSGCs in the nasal and ventral domains of *Ptprk* null retinas. At P0, the Superior oDSGC number matched littermate controls (Fig. 2A-B); however, by P8 we observed a 50% reduction: an average 30.98 cells/mm^2^ (SD = 4.15) in *Ptprk* mutant retinas vs. 60.64 cells/mm^2^ (SD = 2.47) in controls (Fig. 2C, D). Notably, *Ptprk*-heterozygous retinas had ∼25% fewer Superior oDSGCs (mean = 45.94 cells/mm^2^; SD = 2.74), indicating a *Ptprk* gene dosage-dependent effect in regulating the number of Superior oDSGCs. In line with the ∼50% reduction in Superior oDSGC number, the Voronoi domain area increased by 2 fold, from an average of 16722.92 µm^2^ (SD = 1221.05) in wildtype to 35363.21 µm^2^ (SD = 1538.96) in *Ptprk* null mutants (Fig. 2E). However, the Voronoi domain regulatory index (VDRI), a measure of cell spacing^23^, remained the same as in littermate controls in *Ptprk* heterozygous and homozygous mice; all three genotypes had an average VDRI of 3, significantly higher than that of a simulated random distribution (Fig. 2F). Thus, Superior oDSGCs in both *Ptprk* heterozygous and null retinas maintained regular mosaic spacing of Superior oDSGCs, suggesting that mechanisms governing oDSGC mosaic patterning remain intact and do not depend on *Ptprk*.

**Figure 2.**
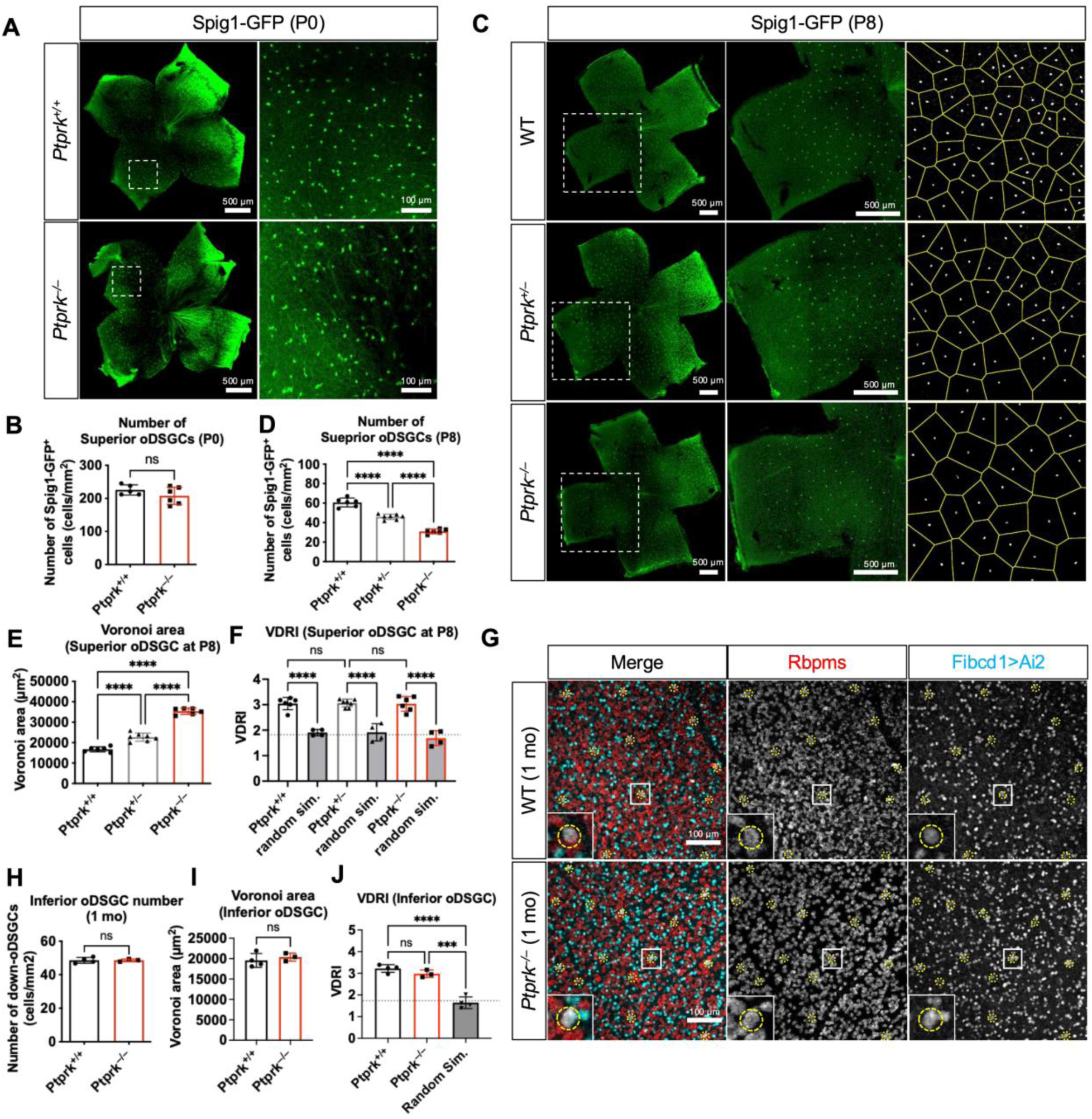
Loss of *Ptprk* leads to a reduction in Superior, but not in Inferior, oDSGC numbers. **(A, B)** The number of Superior *Spig1-GFP^+^* cells, or Superior oDSGCs, in *Ptprk^−/−^* remain comparable to the control at P0. **(C, D)** At P8, both *Ptprk^+/–^* and *Ptprk^−/−^* retinas have reduced numbers of *Spig1-GFP^+^* cells. Examples of Voronoi mosaic analysis using *Spig1-GFP^+^* cells are shown in (C) in the right column. **(E, F)** Voronoi area (E) was quantified and used to calculate Voronai domain regularity index (VDRI) (F), which equals the average of Voronoi area divided by SD. The VDRI of each genotype was compared to that of random simulated mosaics (random sim.) of the same cell density. The dashed grey line indicates the VDRI of a random array of points (1.91) ^23,40^. (G) Inferior oDSGCs in 1 month old (mo) retinas were identified via *Fibcd1^Cre^*-driven expression of YFP (Fibcd1>Ai2; cyan) combined with Rbpms immunostaining (red). Quantification of the number (H), Voronoi domain (I) and VDRI (J) of Inferior oDSGCs in both *Ptprk^+/+^* and *Ptprk^−/−^*. Data are shown in mean ± SD. N = number of animals, each containing two retinas. \*\*\**p* < 0.001, \*\*\*\**p* < 0.0001.

We next asked whether Inferior oDSGCs were also affected by loss of *Ptprk*. Using a *Fibcd1^Cre^* driver mouse line to label Inferior oDSGCs based on the selective expression of *Fibcd1* in Inferior oDSGCs (Fig. S2A-C) ^15^, we detected no change in Inferior oDSGC number (Fig. 2G, H) or mosaic spacing (Fig. 2I, J) at 1 month of age in *Ptprk* mutants, indicating that *Ptprk* functions to preserve postnatal Superior, but not Inferior, oDSGC number. Taken together, these results show that *Ptprk* contributes to the survival of many Superior oDSGCs in early postnatal development but is not required for the survival of ∼50% of these oDSGCs. Further, we find that *Ptprk* maintains the density of Superior, but not Inferior oDSGCs, in a dose-dependent manner.

### *Ptprk* is required for normal Superior oDSGC dendritic morphology

Since neuronal morphology, in particular dendritic architecture, critically influences neural circuit connectivity and ultimately neural computations^24,25^, we next examined the dendritic arbors of the remaining Superior oDSGCs in *Ptprk* mutant retinas. Although the Superior-oDSGC number is reduced in these mutants, their dendrites maintain normal laminar stratification in the adult retina, arborizing primarily within the S4 sublamina with only a very minor presence in S2; this is a pattern identical to that observed in control retinas^14^ (Fig. 3A, B). Examination of retinal cross sections revealed a notable reduction in the thickness of *Ptprk^−/−^* global null retinas, which was also observed following *Chx10^Cre^; Ptprk^f/f^* pan-retinal loss-of-function (LOF). This reduction in retina thickness includes a loss within the IPL of processes within sublamina S3, as indicated by a total loss of S3 Calbindin immunostaining (Fig. S3A) and an overall reduction in the thickness of the IPL and the INL (Fig. S3B, C, F, G). Both global and pan-retinal *Ptprk* loss-of-function retinas exhibit IPL thinning between sublaminae S2 and S4, as indicated by ChAT immunostaining (Fig. S3D, E). Notably, no changes were observed in the thickness of the photoreceptor layer. These observations suggest that cell loss in INL leads to select IPL innervation deficits. However, key AOS circuitry components that are required for the OKR response (vGlut3^+^ amacrine cell processes, ChAT^+^ SAC dendrites; Fig. S3A and SAC number (Fig. S3H, I)), remain intact following *Ptprk* loss of function.

**Figure 3.**
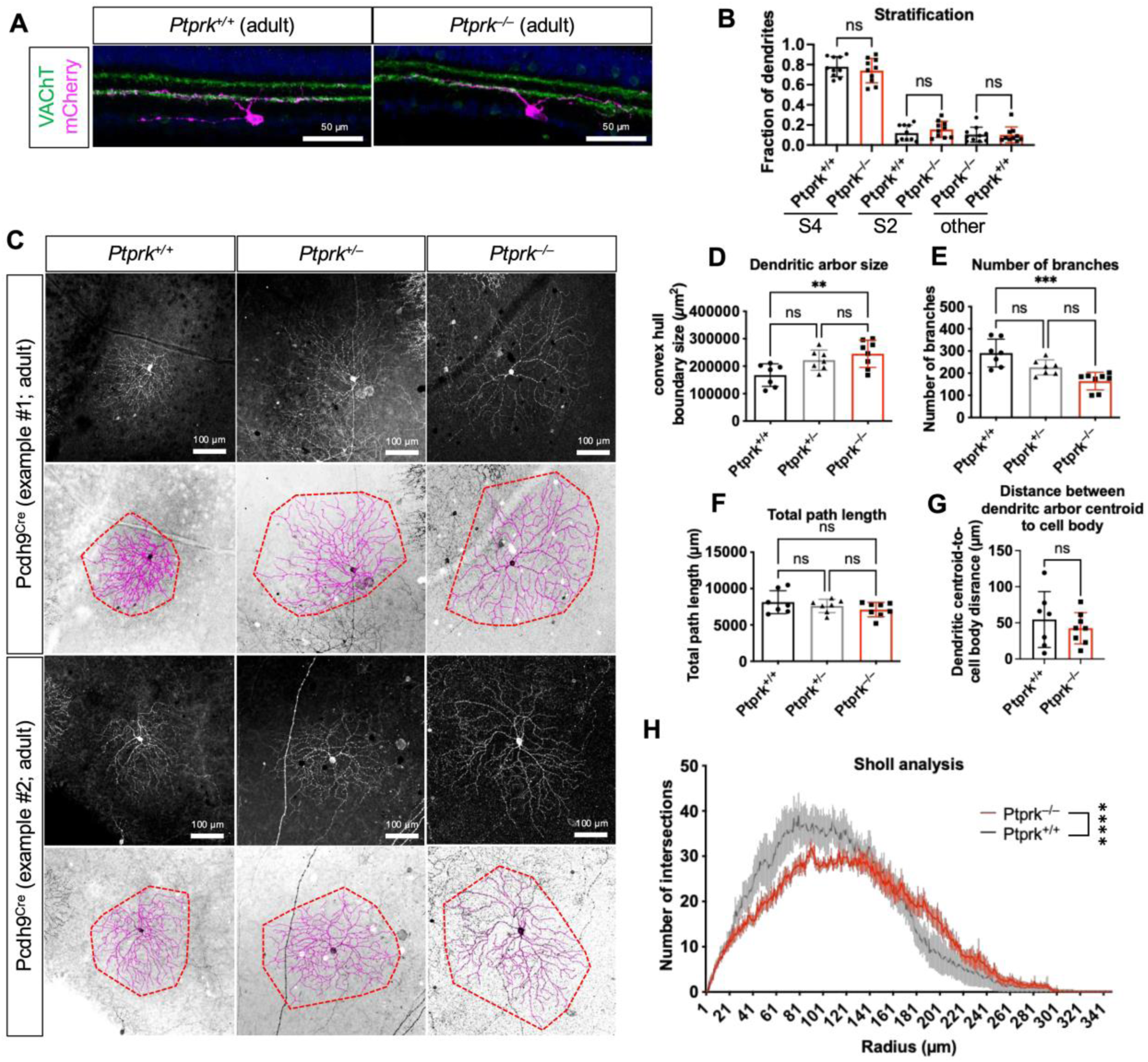
*Ptprk*-null Superior oDSGCs exhibit altered dendritic morphology. **(A, B)** Cross-sections of adult retinas in which Superior oDSGCs were labeled using *Pcdh9^Cre^* and AAV-based brainbow (BChT) virus to express Cre-dependent mCherry. The fraction of the dendrites that colocalized with layer S4, S2 (marked via VAChT) or not (other) were quantified (B). Both *Ptprk^−/−^* and the control had a predominant stratification at S4. **(C)** Superior oDSGC dendrites in *Ptprk^+/+^, Ptprk^+/–^* and *Ptprk^−/−^* retinas were labeled with mCherry using *Pcdh9^Cre^* combined with the BChT virus used in (A) to express mCherry. This approach labeled both Superior oDSGCs and SACs^26^ and thus only the Superior oDSGCs that were isolated from other labeled cells and were neither at the periphery nor in the center of the retina were traced for dendritic morphology analysis. Two example cells for each genotype were presented here and the dendritic arbor areas were delineated via red dashed lines. **(D-F)** Analysis of dendritic traces showed that *Ptprk^−/−^* exhibited an increased in dendritic arbor size (D) and decreased number of dendritic branches (E) while the total path length of all dendrites of a cell remained unchanged compared to the control (F), leading to a rightward shift on the Sholl analysis plot (H). **(G)** The asymmetry of the dendritic orientation was quantified via calculating the distance between the soma to the geometric center, or centroid of the dendritic arbor; the longer the distance, the more asymmetric the dendritic orientation is. No difference in dendritic orientation asymmetry was detected between the null and the control. Data are shown in mean ± SD. N = number of cells. \*\**p* < 0.01, \*\*\**p* < 0.001, \*\*\*\**p* < 0.0001.

Though Superior oDSGCc dendritic laminar stratification was unchanged, we found in *Ptprk^−/−^* mutants significant changes in dendritic morphology. We labeled Superior oDSGC dendrites using *Pcdh9^Cre^*-driven expression of mCherry, delivered via sparse AAV infection of adult *Ptprk^−/−^* and wild-type retinas^26^ (Fig. 3C). We observed that dendritic arbor areas were expanded ∼1.6-fold: from an average 167564.59 µm^2^ (SD = 4137.26) in controls to 245202.43 µm^2^ (SD = 49536.77) in *Ptprk* null retinas, commensurate with the increase in Voronoi domain area we observed in these same mutants (Fig. 3D). Interestingly, despite increased dendritic field size, the total dendritic path length was unchanged owing to a decrease in the number of dendritic branches in *Ptprk* mutants (Fig. 3E, F). Consistent with the observed reduction in Superior oDSGC cell number in *Ptprk* mutants, *Ptprk^+/–^* Superior oDSGCs exhibit an intermediate dendritic expansion phenotype, highlighting the *Ptprk* dosage-dependent role in regulating both cell density and dendritic morphology (Fig. 3C-F). This change in dendritic morphology led to a rightward shift in the Sholl analysis plot (Fig. 3H), reflecting this change in dendritic complexity. However, *Ptprk* LOF does not appear to affect dendritic arbor symmetry since the distances between Superior oDSGC cell bodies and the geometric center of the dendritic arbor (centroid) was unchanged in *Ptprk* null retinas (Fig. 3G). These results show that Superior oDSGC density and dendritic arbor size are correlated, suggesting compensatory remodeling of oDSGC dendritic architecture in response to reduced cell density resulting from loss of PTPRκ.

### Loss of *Ptprk* increases spike responses and broadens oDSGC tuning curves

Since *Ptprk* influences Superior oDSGC cell number and dendritic morphology, we next asked if *Ptprk* affects the physiological properties of vertically-tuned oDSGCs. To identify vertically-tuned oDSGCs, we injected retrograde tracers (Retrobeads) into the MTN and targeted fluorescently-labeled cell bodies via two-photon imaging in isolated retinas^27^. We generated tuning curves of oDSGCs by measuring extracellular spikes in response to moving bar stimuli in 8 directions (Fig. 4A). Both Superior and Inferior oDSGCs were present in control and *Ptprk^−/−^* retinas (Fig 4B), defined by their preference for ventral (Superior) or dorsal (Inferior) motion across the retina. In *Ptprk* null mice, oDSGCs generated more spikes in response to stimuli moving in all directions, indicated by larger tuning curve areas (Fig 4C) compared to controls; however, both *Ptprk* null and control oDSGCs exhibited fewer spikes in their preferred direction (Fig. 4D). We used two metrics to assess oDSGC tuning: (1) the shape of the tuning curves when normalized by the maximum spike response (normalized tuning curve area) and (2) the direction selectivity index (DSI) defined as the vector sum divided by the scalar sum of responses. Normalized tuning curve areas were larger (Fig. 4E) and direction selectivity indices were lower (Fig. 4F) in *Ptprk* null mutants compared to controls. This indicates that loss of *Ptprk* results in more broadly tuned, less direction selective, vertically-tuned oDSGCs. These changes in tuning curves are consistent with oDSGCs spiking more in non-preferred directions, while preferred direction responses remain unchanged, resulting in diminished direction selectivity. These results demonstrate that loss of *Ptprk* impacts oDSGC function by increasing spike responses and reducing direction selectivity in the retina.

**Figure 4.**
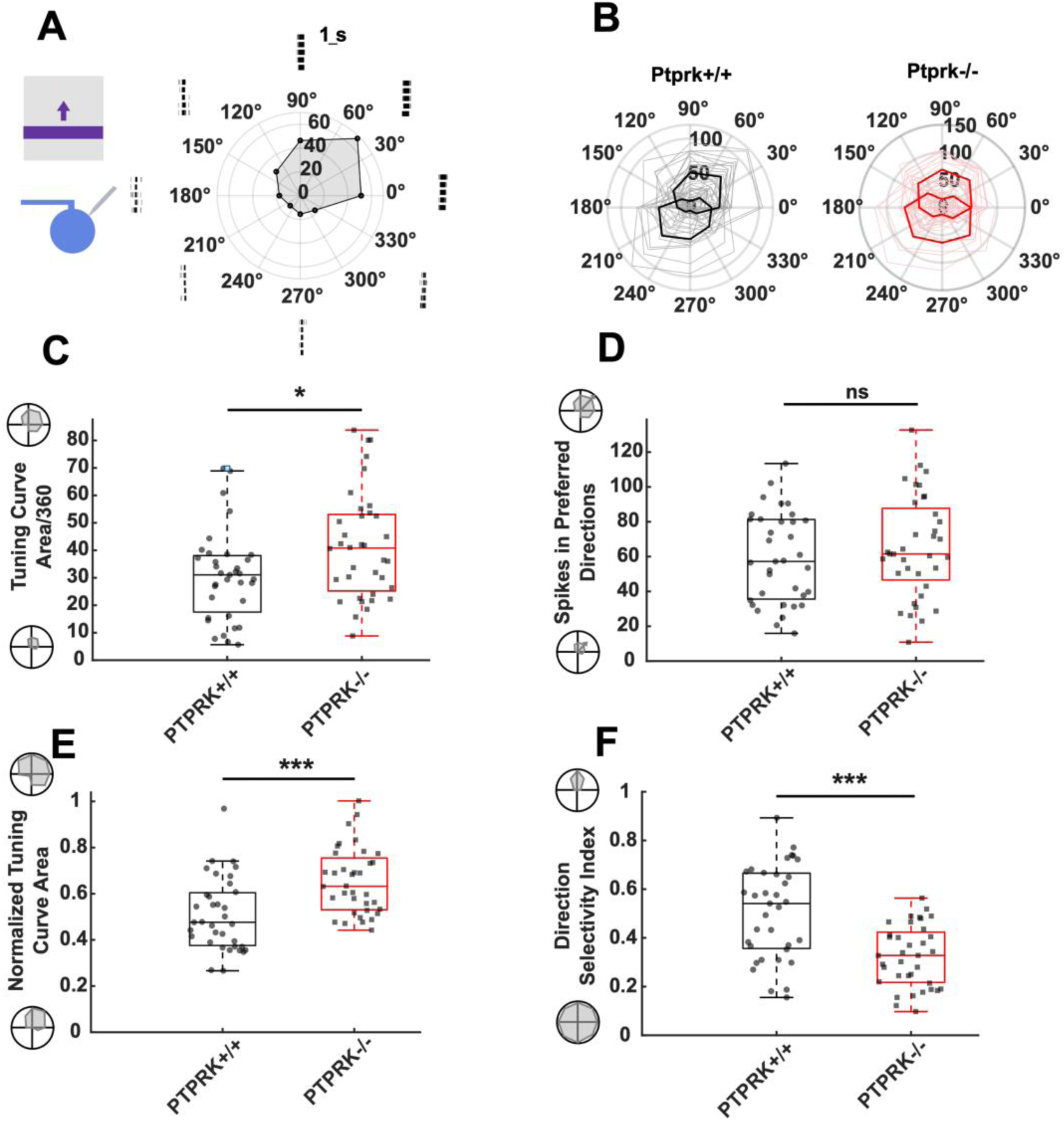
*Ptprk*-null oDSGCs exhibit broader tuning curves. **(A)** Direction selectivity of a wildtype Inferior oDSGC is shown in retinal space as the mean tuning curve of spike responses to moving bars in 8 directions. Raster plots indicate responses to each direction across five trials. **(B)** Tuning curves for oDSGCs in control (black) and *Ptprk*−/− (red) retinas demonstrate presence of cells with preference for two directions,Superior (30-150°) and Inferior (210-330°) cell types, shown as mean tuning curves for each direction (thicker lines). **(C-F)** Comparisons of (C) tuning curve area (p=0.018), (D) the number of spikes in the cell’s preferred direction (p=0.41), (E) the normalized tuning curve area (p<0.001) **(F)** and direction selectivity index (p<0.001), indicate that *Ptprk^−/−^* oDSGCs are more broadly tuned than control. (Control (black): N=35; *Ptprk^−/−^* (red): N=36).

### Loss of *Ptprk* disrupts Superior oDSGC direction selectivity

We next evaluated changes in Superior and Inferior oDSGCs tuning in the context of *Ptprk* loss of function. In control and *Ptprk* null retinas, we identified both Superior and Inferior cells, based on their preferred directions (Fig. 4B); however, we found a broader distribution of preferred directions in the *Ptprk* null mutants (Fig. 5A-B). In these mutants we found a subset of MTN-projecting cells with abnormal tuning; these oDSGCs preferred horizontal motion, outside of the known preferred directions of Superior or Inferior cells (Fig. 5C). Among littermate controls, such horizontal tuning was seldom observed (Fig. 5B). These results show that *Ptprk* influences the directional preference of a population of MTN-projecting oDSGCs.

**Figure 5.**
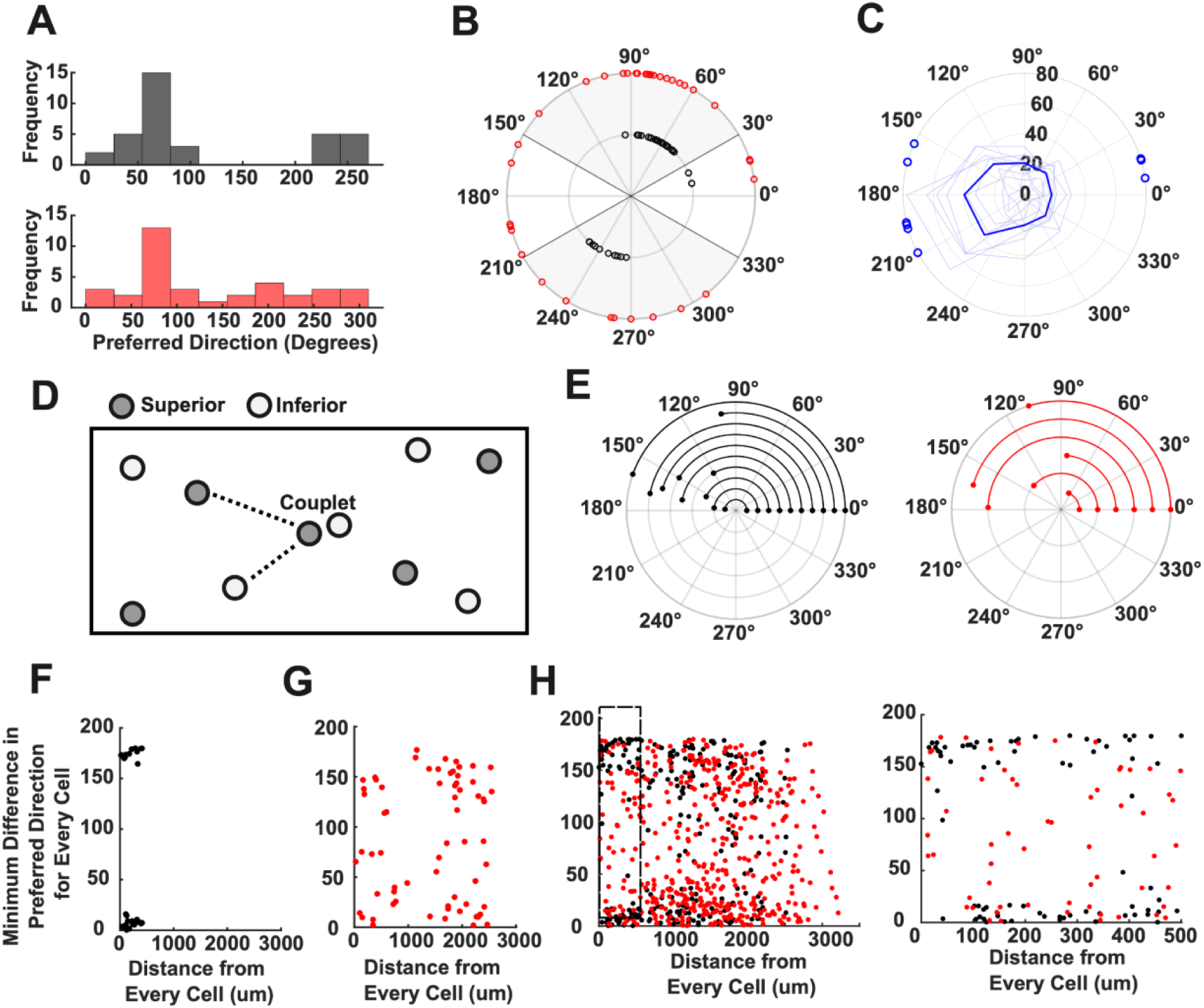
*Ptprk*-null oDSGCs have a broader distribution of preferred directions. **(A)** Histograms of oDSGC preferred directions demonstrate two distinct peaks in control (black, top) and a broader distribution in *Ptprk^−/−^* (red, bottom) retinas.**(B)** Polar coordinate representation of preferred directions where grey sectors indicate preferred direction selection criteria for Inferior (30-150°) and Superior (210-330°) oDSGCs based on previous wildtype datasets^27^. **(C)** Population of *Ptprk^−/−^* oDSGCs that are abnormally tuned, with preferred directions that did not fall into Superior and Inferior classifications**. (D)** Diagram of oDSGC mosaics represents proximity of oDSGC couplets and nearest neighbor distances to same or opposing cell types (Superior: dark gray, Inferior: light gray). **(E)** For oDSGC couplets within 30µm of each other, the minimum difference in preferred directions shown in polar coordinates is larger in control (black, left) compared to couplets identified in *Ptprk^−/−^* (red, right). **(F-G)** Comparison between the distance from every cell and the minimum difference in preferred direction is shown for a representative retina in (F) control and in (G) *Ptprk^−/−^*, demonstrating two distinct populations in control retinas which are absent in *Ptprk^−/−^*. **(H)** The same comparison of the minimum difference between preferred direction as a function of the distance between cell positions for both genotypes across all retinas (left) and for a distance less than 500 µm (right) indicates greater variability in *Ptprk^−/−^*oDSGCs. (Control (black): Superior N=12, Inferior N=23; *Ptprk*^−/−^ (red): Superior N=8, Inferior N=18, Abnormally tuned N=8).

Next, we asked if the effect of *Ptprk* LOF on oDSGC tuning is specific to oDSGC type. Because our method of discriminating between Superior or Inferior cell identity relies on the cell’s preferred direction, we used the spatial distribution of oDSGC types to discriminate between the cell types. Since Superior and Inferior oDSGCs form independent mosaics, the spatial distribution of each type ensures that two neighboring cells of the same type have a larger nearest neighbor distance than those of opposing types (Fig. 5D). This results in the closest neighboring oDSGCs (< 30 µm, called “couplets”) being distinct types and having a difference of approximately 160° in their preferred directions. We first observed an overall ∼25% decrease in the total population of vertically tuned oDSGCs in both *Ptprk* null (−33%) and pan retinal mutants (−20%), an outcome one would predict with 50% loss of Superior oDSGCs and unchanged Inferior oDSGCs (Fig. S4A-C, E). Although this led to a significant decrease in the number of couplets (Fig. S4D, F), we were still able to examine preferred directional tuning within oDSGC couplets (Fig 5E). oDSGC couplets in control retinas had opposing preferred directions, resulting in an average minimum difference in their preferred directions of 153.08° (range: 98.58-173.09°) (Fig 5F). In contrast, we observed that the preferred directions of oDSGCs in *Ptprk* knockout couplets have a smaller average minimum difference in preferred directions (122.92°) (range: 65.36-178.33°). Notably, at least one cell in each couplet had a preferred direction that fell within the Inferior oDSGC classification: a preferred direction between 30° and 120°, suggesting that in couplets, abnormal tuning often occurs in the non-Inferior oDSGC partner, the Superior oDSGC (Fig. 5E, G). Thus, we call this abnormally-tuned ganglion cell an incorrectly tuned Superior cell. Given that Superior and Inferior oDSGCs each maintain their mosaics (Fig. 2), we expect that neighboring oDSGCs of opposing types will be closer than those of the same type. Therefore, we compared the distance between oDSGCs with the minimum difference in their preferred directions and found different patterns comparing control and *Ptprk* knockout oDSGCs (Fig. 5F-H). In control retinas, the preferred directions were either similar to the reference cell (likely the same type of oDSGC) or displayed approximately a 160° difference (likely the opposing type), and oDSGCs of the same type rarely occurred within 100 µm of the reference cell (Fig 5H). In *Ptprk* knockouts, however, oDSGCs had preferred directions within 160 degrees of the reference cell, and the same type of oDSGC often occurred close to the reference cell (within 100 µm) (Fig. 5H). These results indicate that the abnormally-tuned direction selective ganglion cells are most likely Superior oDSGCs situated next to normally-tuned Inferior oDSGCs. Despite being aberrantly and sometimes horizontally tuned, these Superior oDSGCs project exclusively to the MTN rather than the NOT (Fig. S4G). Therefore, *Ptprk* LOF causes a disruption of Superior oDSGC direction selectivity without affecting their central projection.

### Loss of *Ptprk* decreases dorsal medial terminal nucleus innervation and also post-synaptic Superior oDSGC connectivity

Does the ∼50% loss of Superior oDSGCs impact post-synaptic connections of the remaining Superior oDSGCs? To answer this question, we intravitreally injected fluorophore-conjugated cholera toxin subunit B (CTB), labeling all retinofugal axons and their retinorecipient targets. Using brain clearing and light sheet microscopy, we observed that the overall MTN morphology, including its ventral-to-dorsal extension, was not affected following either global or the pan-retinal *Ptprk* LOF (Fig. 6A-B). However, though oDSGC projections to vMTN were preserved, the innervation of dMTN, the primary target of up-oDSGCs, was significantly attenuated, as evidenced by reduced area of labeled RGC axons in the dMTN (Fig. 6A-E). No comparable alterations in RGC innervation were observed in other retinorecipient targets, including the olivary pretectal nucleus (OPN), suprachiasmatic nucleus (SCN), or dorsal lateral geniculate nucleus (dLGN) (Fig. S5A-D).

**Figure 6.**
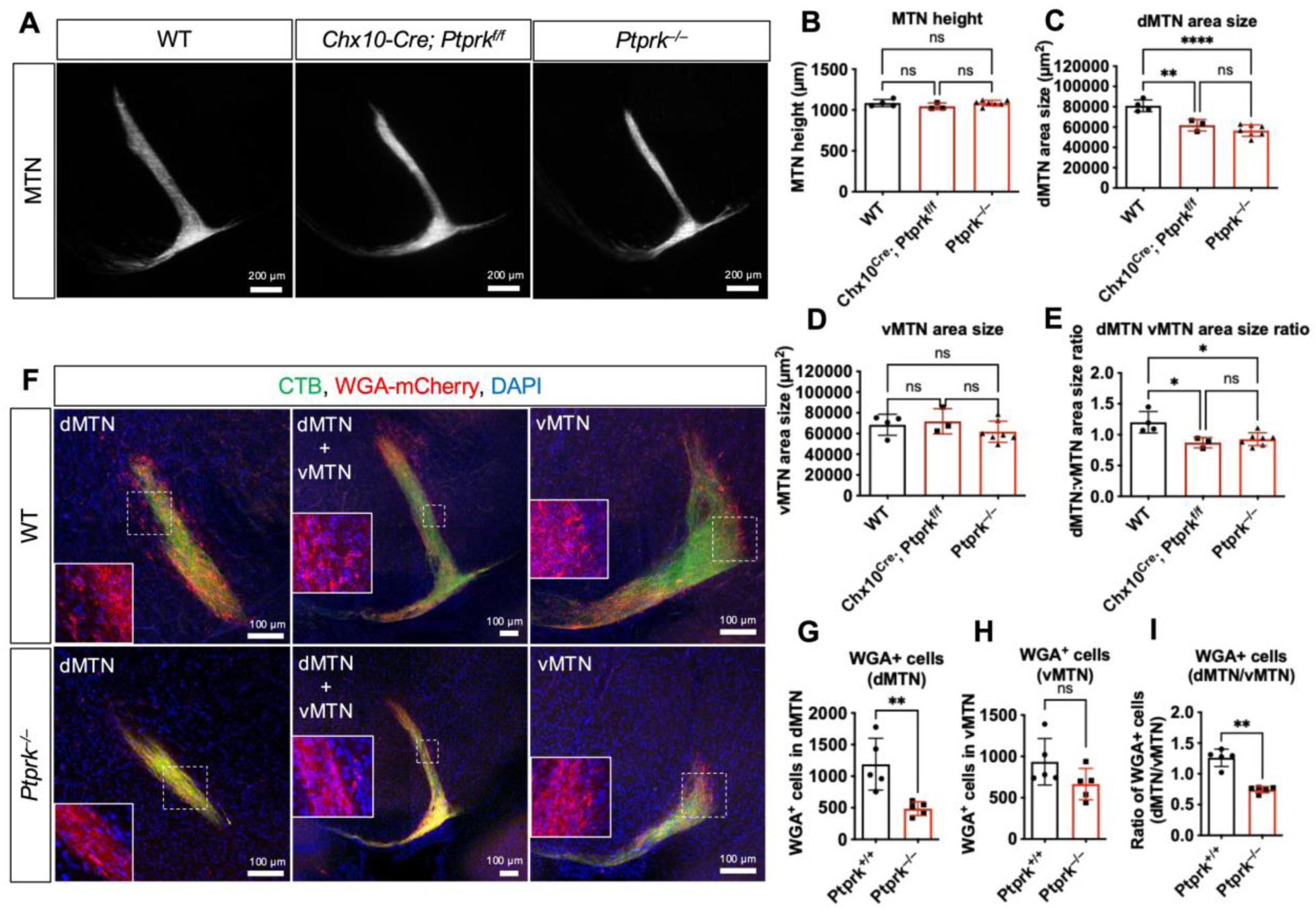
*Ptprk* deletion causes decreased dMTN innervation and fewer connections with post-synaptic neurons. **(A)** Maximum intensity projections of light-sheet images of MTN innervated by RGC axons, labeled via intravitreal injection of CTB. **(B-E)** Quantification of projected 2D images for MTN height (B) and area of the dMTN (C), vMTN (D), and ratio of areas between dMTN and vMTN (E). Since the MTN height remained the same compared to WT, an arbitrary cutoff of 150 µm from the bottom of MTN is used to define dorsal vs ventral MTN. N = number of animals. **(F)** Intravitreal injection of both CTB (green) and AAV-based WGA-mCherry (red) was performed to delineate MTN and anterogradely label MTN neurons postsynaptic of vertical oDSGCs, respectively. **(G, H)** Quantification revealed a significant decrease in WGA^+^ cell number in dMTN, from 1188.6 cells (SD = 409.9) in controls to 486.6 cells (SD = 109.2) in nulls (G), but not in vMTN, with 934.2 cells (SD = 280.7) in controls and 664.8 cells (SD = 189.31) (H). **(I)** A decrease in the ratio of WGA^+^ cell number between dMTN and vMTN in *Ptprk* nulls, from an average 1.29 (SD = 0.14) in controls to 0.74 (SD = 0.05) in *Ptprk^−/−^*. N = number of MTNs. Data are shown in mean ± SD. \**p* < 0.05, \*\**p* < 0.01.

To assess further whether diminished dMTN innervation area is accompanied by a deficit in postsynaptic connectivity, we performed anterograde tracing using an AAV-based fluorophore-conjugated wheat germ agglutinin (WGA) virus to label vertical oDSGC postsynaptic target cells in the MTN (Fig. 6F). Quantification revealed that the number of WGA^+^ postsynaptic cells in the dMTN, but not vMTN, was reduced from an average of 1188.6 (SD = 409.92) cells to 486.6 (SD = 109.21) cells in the entire dMTN, a 59% reduction similar to the reduction in Superior oDSGC number resulting from *Ptprk* loss of function (Fig. 6F-H). This leads to a decrease in the ratio of WGA^+^ cells between dMTN and vMTN: from ∼1.25 to 0.75 (Fig. 6I). Together, these findings show that *Ptprk* loss of function selectively disrupts retinorecipient connectivity to the dMTN, but not the vMTN or other retinorecipient targets, consistent with the reduction we observe in Superior oDSGC number in *Ptprk* mutants.

### Loss of *Ptprk* disrupts the vertical OKR via a retinal mechanism

To assess whether the reduction in Superior oDSGC population affects AOS image stabilization, we recorded OKR responses from *Ptprk* LOF mice. Though the null mice exhibited normal horizontal tracking in response to continuously drifting stimuli, their vertical OKR performance was significantly attenuated in both ventrodorsal (superior) and dorsoventral (inferior) directions (Fig. 7A-E). Assessment of eye tracking in response to sinusoidally moving stimuli also revealed vertical, but not horizontal, response deficits in the OKR gain, reflected in eye movements in response to the moving stimulus (Fig 7F, G).

**Figure 7.**
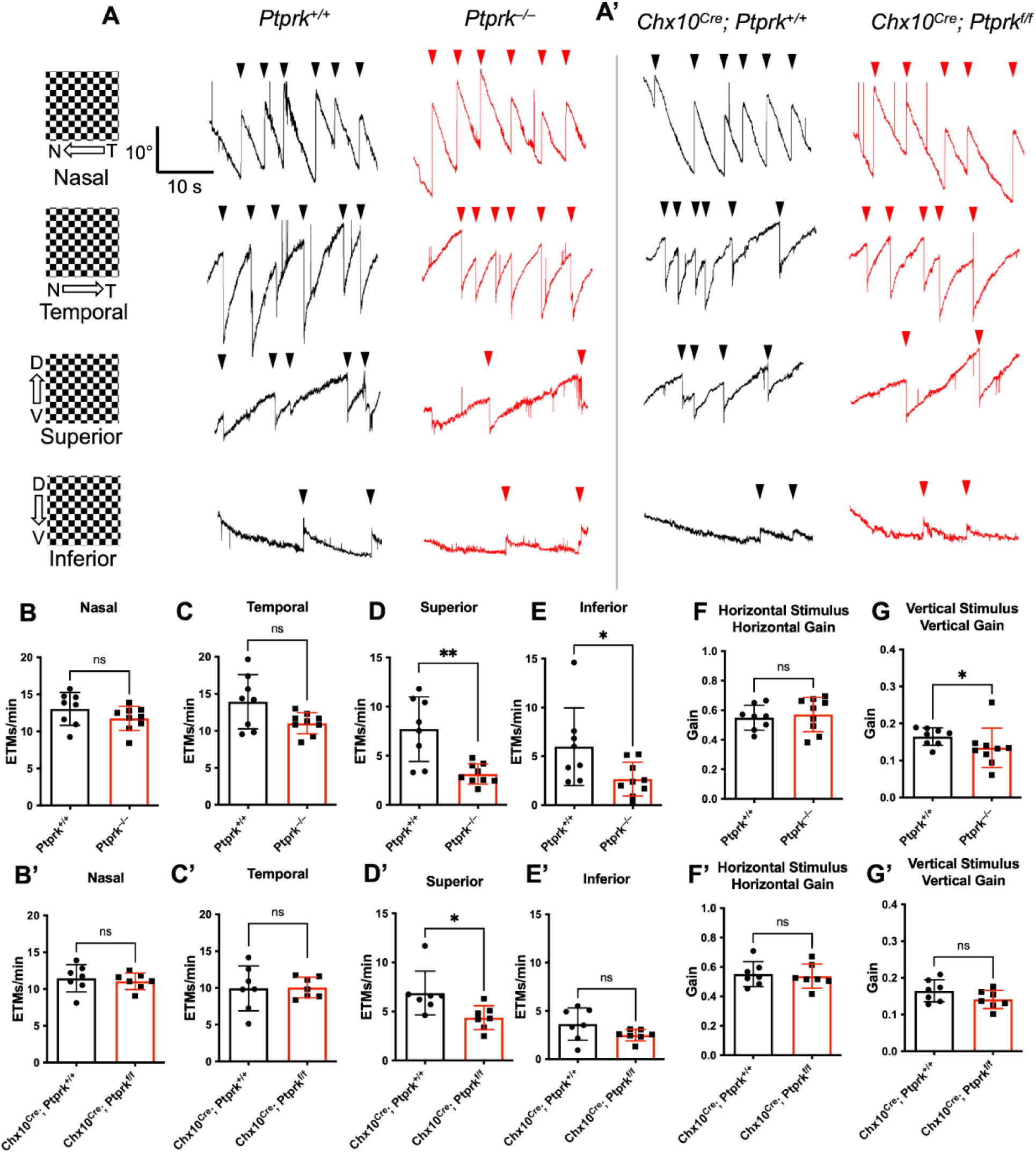
Superior OKR is attenuated in both global and pan-retinal *Ptprk* knockouts. **(A-A’)** OKR responses of adult mice from (A) *Ptprk^+/+^*, *Ptprk^−/−^* (A’) *Chx10^Cre^; Ptprk^+/+^* and *Chx10^Cre^; Ptprk^f/f^* in nasal, temporal, Superior and Inferior directions. **(B-E, B’-E’)** Quantification of eye tracking movements (ETMs), each containing a slow object tracking phase along the direction of motion and a fast compensatory saccade in the opposite direction (indicated by arrowheads in A-A’), for (B) Nasal, (C) Temporal, (D) Superior, and (E) Inferior stimuli. **(F, G, F’, G’)** Quantification of eye-tracking gains of (F) horizontal and (G) vertical axes, with 0 indicating no tracking and 1 perfect tracking. Data are shown in mean ± SD. N = number of animals. \**p* < 0.05, \*\**p* < 0.01.

To determine if the *Ptprk*-dependent behavioral defect is driven by a retinal mechanism, we generated a *Ptprk* conditional allele and crossed it with a pan-retinal Cre driver line, *Chx10^Cre^*. Mice lacking *Ptprk* in the retina exhibited similar superior OKR defects as in the global *Ptprk* null in response to superior drifting gratings (Fig. 7A’-D’), but no significant changes to the inferior drifting gratings (Fig 7E’) or to the sinusoidally drifting gratings (Fig. 7F’-G’). This difference could reflect incomplete removal of *Ptprk* by the *Chx10^Cre^* allele. Aside from these vertical OKR deficits, both the global and conditional null animals exhibited normal pupil constriction (Fig. S6A), normal spontaneous saccades (Fig. S6B), and normal fear responses to looming threats (Fig. S6C). These results show that retinal *Ptprk* contributes to the amplitude of vertical, in particular superior, eye movements that define the OKR.

## Discussion

There are more than 45 types of RGCs, and most form a mosaic with different densities, depending on the cell size and coverage factor (overlap between dendrites). However, the mechanisms by which each RGC type is specified and maintained at the correct abundance—ensuring appropriate matching of synaptic partners and functional circuit assembly—remain poorly understood. Here, we identify a new role for the protein tyrosine phosphatase receptor type kappa (Ptprk) in maintaining the number and tuning properties of Superior oDSGCs, an essential component of the accessory optic system that drives the OKR. High-depth scRNAseq and RNAscope *in situ* hybridization revealed that *Ptprk* is enriched in Superior relative to Inferior oDSGCs, a result supported by multiple published scRNAseq datasets. Loss of *Ptprk* reduced Superior, but not Inferior oDSGC number by P8. Despite this oDSGC loss, the remaining Superior oDSGCs preserve a regular mosaic distribution, resulting in expanded intercellular spacing and expansion of the Superior oDSGC dendritic arbors, likely reflecting a compensatory mechanism to preserve dendritic coverage of the retina and, hence, the Superior oDSGC mosaic. *Ptprk* loss also leads to decreased oDSGC innervation of dMTN, diminishing connectivity with downstream post-synaptic partners, presumably caused by decreased Superior oDSGC number. Physiologically, *Ptprk* LOF broadened tuning in oDSGCs and generated aberrant directional tuning in about half of Superior oDSGCs. Lastly, we analyzed OKR performance using both a *Ptprk* global null allele and a newly generated conditional allele. In both global and pan-retinal knockouts, superior OKR responses were selectively attenuated, suggesting that *Ptprk* within the retina regulates superior OKR. Together, these findings show that *Ptprk* loss reduces Superior oDSGC density, causes aberrant tuning in the remaining Superior oDSGCs, weakens dMTN activation, and consequently limits MTN drive and superior OKR responses. These results highlight the importance of maintaining the correct abundance in a cell type-specific fashion, demonstrating how decreased neuronal density can lead to neurite changes and, ultimately, to a reshaping of circuit function and neural computation.

We found that Superior oDSGC numbers were normal at P0 but reduced by 50% at P8. This timing coincides with the developmental wave of RGC apoptosis, during which approximately half of all RGCs present at birth are eliminated by P5^28,29^. Since Superior oDSGCs are generated in normal numbers at P0, their subsequent loss in *Ptprk* mutants likely reflects increased programmed cell death rather than defective specification. Although *Ptprk* is well known for its tumor-suppressive roles, it also modulates cell-death pathways and can promote neuronal survival^30^. *Ptprk* knockdown has been shown to elevate caspase-3, caspase-8, and p53 expression, as well as increasing phosphorylated JNK levels: these effects can collectively enhance apoptosis in prostate cancer models^30,31^. One key substrate of PTPRκ is the epidermal growth factor receptor (EGFR); cells with elevated EGFR levels undergo apoptosis upon EGF stimulation, and PTPRκ-mediated dephosphorylation of EGFR can limit hyperactivation and attenuate apoptotic stress^32–34^. PTPRκ can also dephosphorylate and inhibit RAS, thereby reducing ERK signaling, excessive activation of which is known to induce apoptosis^35^. In addition, activated PTPRκ undergoes proteolytic cleavage by Furin, ADAM10, and γ-secretase, generating an intracellular domain that translocates to the nucleus to enhance β-catenin-dependent transcription through LEF/TCF^36^. β-catenin activation promotes the release of trophic factors such as Lif and Fgf2, which support retinal neuron survival through a STAT3-dependent mechanism^37^. This suggests that the cleaved extracellular domain of PTPRκ may serve paracrine-like functions for the survival of neighboring cells. Since *Ptprk* exhibits a gene dosage effect with respect to regulating Superior oDSGC number, the level of a secreted PTPRκ extracellular domain in the retina could be a determining factor for Superior oDSGC density. Together, these signaling pathways provide several plausible mechanisms through which PTPRκ protects Superior oDSGCs during the developmental cell-death window. However, the precise downstream signaling events through which *Ptprk* maintains cell number in the retina remain to be elucidated.

Our scRNAseq dataset, along with previous work, provides evidence for genetic differences between oDSGC types. Yet, the molecular mechanisms that determine direction selectivity remain unknown. Here, we report evidence of abnormal tuning in a population of Superior oDSGCs due to the loss of *Ptprk*. *Ptprk* is not expressed in SACs (Fig S1) which provide tuned inhibition to DS circuits, but it is expressed in type 5 bipolar cells^18^ and Superior oDSGCs. Because *Ptprk* is expressed during early development, it is possible that *Ptprk* is involved in matching oDSGC synaptic partners during the development of retinal DS circuits. The selectivity of abnormal tuning to a subset of Superior cells suggests that Ptprk is not necessary for general DS tuning and so additional *Ptprk*-independent factors are likely relevant for these responses to image motion. Future studies that assess synaptic connectivity and excitatory/inhibitory currents of abnormally tuned oDSGCs in *Ptprk* null mutants will help define molecular cues critical for the development and maintenance of DS circuits.

The novel finding here of oDSGCs with aberrant preferred directions and disrupted vertical OKR following the loss of *Ptprk* provides insight into the processing of image motion by the AOS. In the control mice, the difference in spike responses between Superior and Inferior oDSGCs to their preferred directions can predict OKR gain^27^. In the global *Ptprk* null mutants, both Superior and Inferior OKR responses were reduced, yet, oDSGCs generated more spikes. One explanation is that the abnormally tuned oDSGCs in *Ptprk* null mutants contribute additional spikes to vertical directions, changing the balance between Superior and Inferior spikes and, subsequently, the OKR. In addition, because we defined oDSGC type identity by observing preferred direction, abnormal Superior cells may be misidentified as Inferior cells and detect downward rather than upward motion, further disrupting downstream OKR computations. Along with physiological changes in the Superior oDSGC population, the 50% reduction in Superior oDSGCs and also the unequal innervation in the *Ptprk* mutant of the dorsal and ventral MTN may also alter OKR output resulting from an underrepresentation of Superior cell detection of upward motion. The effects of *Ptprk* LOF on Superior oDSGC survival, tuning, and MTN innervation are likely the main drivers of observed OKR deficits. However, further study of MTN function will clarify how loss of *Ptprk*-dependent mechanisms disrupt AOS computations with behavioral consequence.

While the functional role of retinal *Ptprk* in regulating the superior OKR is robust, global *Ptprk* null animals exhibited deficits in the inferior OKR whereas pan-retinal conditional mutants did not show a significant difference in the inferior OKR. Because the conditional allele deletes exon 3, the same lesion present in the global *Ptprk* null mutant, in all Cre-expressing cells, we attribute these phenotypic differences between global and conditional mutants to the efficiency of the *Chx10^Cre^*-mediated recombination. Although *Chx10* is expressed in the vast majority of retinal progenitors during embryogenesis and therefore *Chx10^Cre^* is widely used as a pan-retinal driver. However approximately 5% of retinal neurons are not derived from *Chx10*-expressing progenitors^38^. This interpretation is supported by our result from Retrobead tracing in adults, which revealed a smaller reduction of oDSGCs in the conditional mutants as compared to the global null mutant. Such “escaper” cells that never underwent Cre-mediated *Ptprk* knockout due to lack of *Chx10* expression could partially preserve Superior oDSGC numbers, thereby mitigating the severity of the vertical OKR deficit in the conditional mutants.

The observation that only half of Superior oDSGCs are lost in *Ptprk* mutants suggests the existence of two distinct subpopulations of Superior oDSGCs. One possibility is that all Superior oDSGCs express *Ptprk*, but only a subset solely relies on PTPRκ-dependent pathways for survival. Alternatively, Superior oDSGCs may consist of two molecularly distinct groups, one *Ptprk*⁺ and one *Ptprk*⁻, with the latter maintained through *Ptprk*-independent mechanisms. Additional experimental validation will be required elucidate this heterogeneity. Further, it is possible that *Ptprk* acts stochastically in Superior oDSGCs to reduce cell number of this cell type by half. In addition, although we identified *Ptprk* as a key regulator of cell density in a specific RGC type, other RGC types may employ similar strategies using distinct molecules. This raises the intriguing possibility that different protein tyrosine phosphatase receptors contribute to type-specific RGC survival. Further investigation into this receptor family may reveal a shared framework governing the maintenance of correct RGC abundance.

To our knowledge, this study provides the first demonstration of a genetic mutation that mistunes direction-selective neurons rather than abolishing direction selectivity altogether. While most perturbations disrupt DS tuning so severely that cells and also the animals lose DS responses, *Ptprk* loss instead produces a population of oDSGCs that retain direction selectivity but adopt incorrect and inconsistent preferred directions. This phenotype reveals an intermediate state, 50% Superior oDSGC loss, that has not been previously described and offers new insight into the mechanisms that determine directional tuning. Our findings suggest that changes in RGC density through a *Ptprk*-dependent mechanism reshape dendritic morphology and subsequently perturb the precise wiring between oDSGCs and their SAC partners, connections that are essential for establishing preferred directions. Further characterization of the aberrantly tuned Superior oDSGCs will be critical for understanding the mechanisms underlying these changes.

In summary, we identify here a single molecule, PTPRκ, that regulates the population size of Superior oDSGCs. *Ptprk* LOF results in: (1) a decrease in the number, but not mosaic regularity, of Superior oDSGCs; (2) broader functional tuning curves across oDSGCs; (3) mistuning of a subset of Superior oDSGCs; (4) a decrease in postsynaptic connectivity between oDSGCs and MTN target cells; and (5) disruption of behavioral superior image stabilization. These findings corroborate work demonstrating the relative resilience of mosaic formation in other retina cell types despite changes in cell density^39^. Further, these findings may extend beyond the accessory optic system, suggesting specific molecular regulation of individual neuronal populations that in turn impacts synaptic connectivity and behavior, highlighting how single molecule perturbations can propagate across scales. Overall, this work provides a framework for studying how developmental dysregulation of defined neuronal types contributes to deficits in sensory processing and, potentially, neurological disorders.

## Figure Legends

**Suppl. Figure 1.**
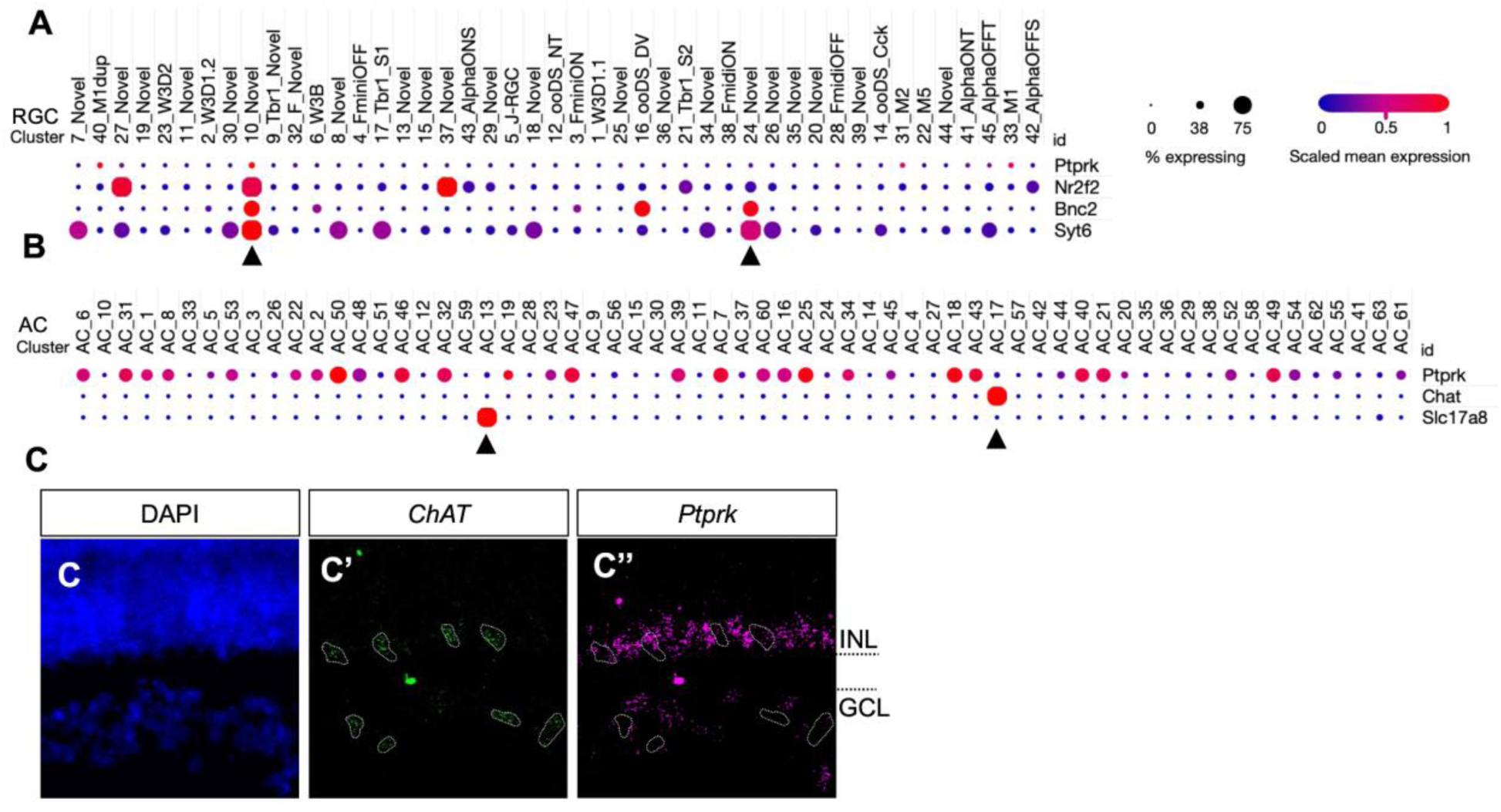
Expression pattern of Ptprk in adult RGCs and neonatal amacrine cells. **(A)** Dotplot generated from P56 RGC dataset by Tran *et al.*^1^ shows that *Ptprk* is expressed in cluster 10 (10_Novel), the putative vertical oDSGC cluster, marked by the expression of *Nr2f2*^20^, *Bnc2*^21^ and *Syt6*^18^, though at a very low level. However, Ptprk is not expressed in cluster 24 (24_Novel) in which putative slow-motion-sensing nasal oDSGCs reside. **(B)** Dotplot generated from P19 amacrine cell datasets^41^ shows that, although *Ptprk* is expressed in various types of amacrine cells, the two types that provide input to the oDSGCs—ChAT-positive SACs and vGlut3 (Slc17a8)-positive ACs—do not express *Ptprk*. **(C)** RNAscope *in situ* hybridization in P8 retina showed that *ChAT* expression does not colocalize with *Ptprk* expression.

**Supple. Figure 2.**
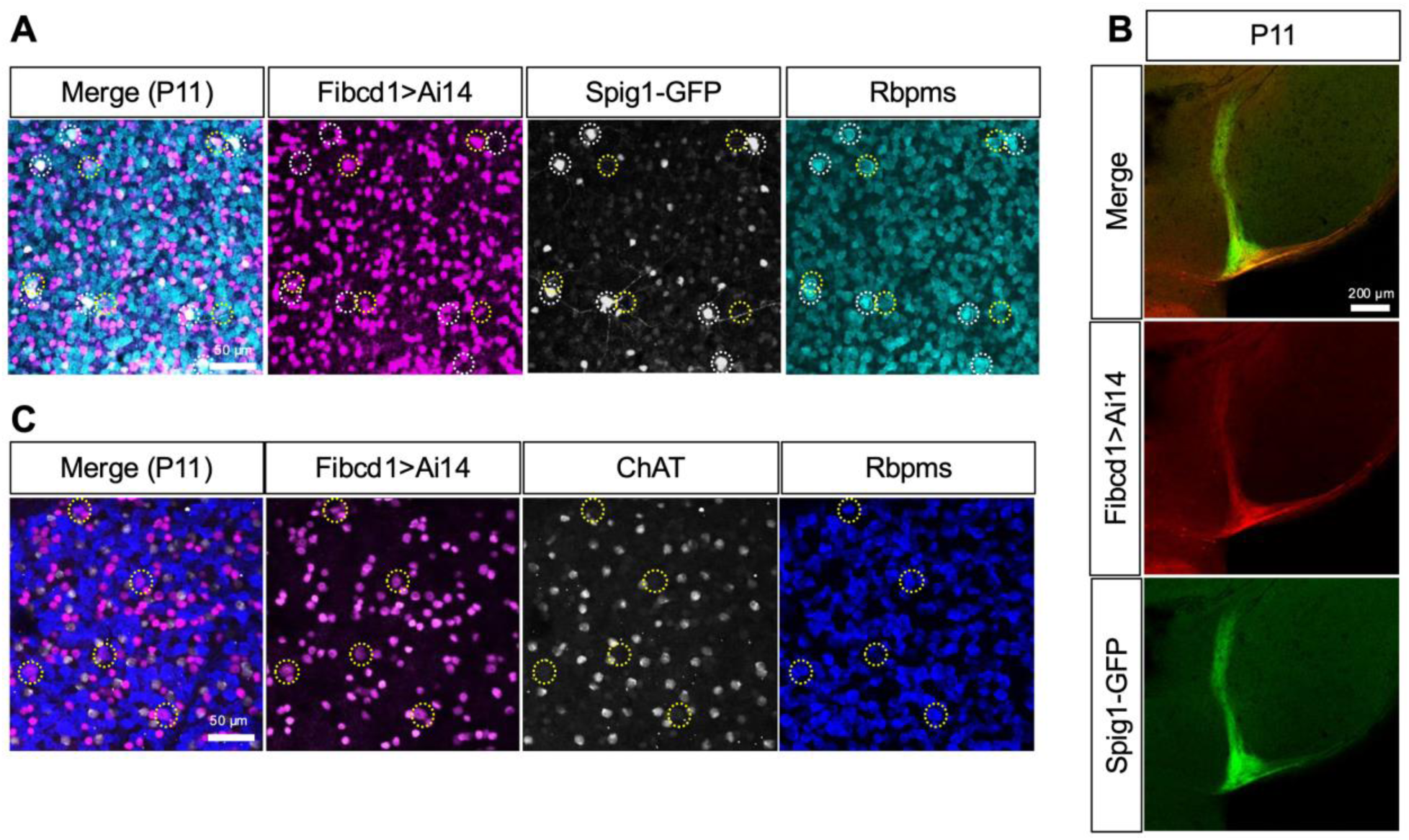
*Fibcd1^Cre^* characterization in Inferior oDSGCs. **(A)** Whole mount retina from P11 mice with *Fibcd1^Cre^*-driven tdTomato expression (Ai14) and Spig1-GFP, a Superior oDSGC reporter. The Cre^+^ RGCs (Rbpms^+^) do not express Spig1-GFP—78/79, i.e., 99% of RGCs across 4 retinas were *Fibcd1^Cre^*-tdTomato^+^ and Spig1-GFP^−^. **(B)** *Fibcd1^Cre^*-tdTomato^+^ RGC axons strongly innervate the ventral portion of the MTN, with the dMTN innervated by Spig1-GFP^+^, suggesting that Fibcd1^+^ RGCs are Inferior oDSGCs. **(C)** Whole-mount P11 retinas show that while there are many Cre-expressing non-RGCs (Rbpms^−^), presumably ACs based on their localization in lower INL and especially GCL, these cells do not express ChAT, excluding them asSACs. Yellow dashed circles indicate Fibcd1^+^ Rbpms^+^ cells; white dashed circles mark Spig1-GFP^+^ Rbpms^+^ cells.

**Supple. Figure 3.**
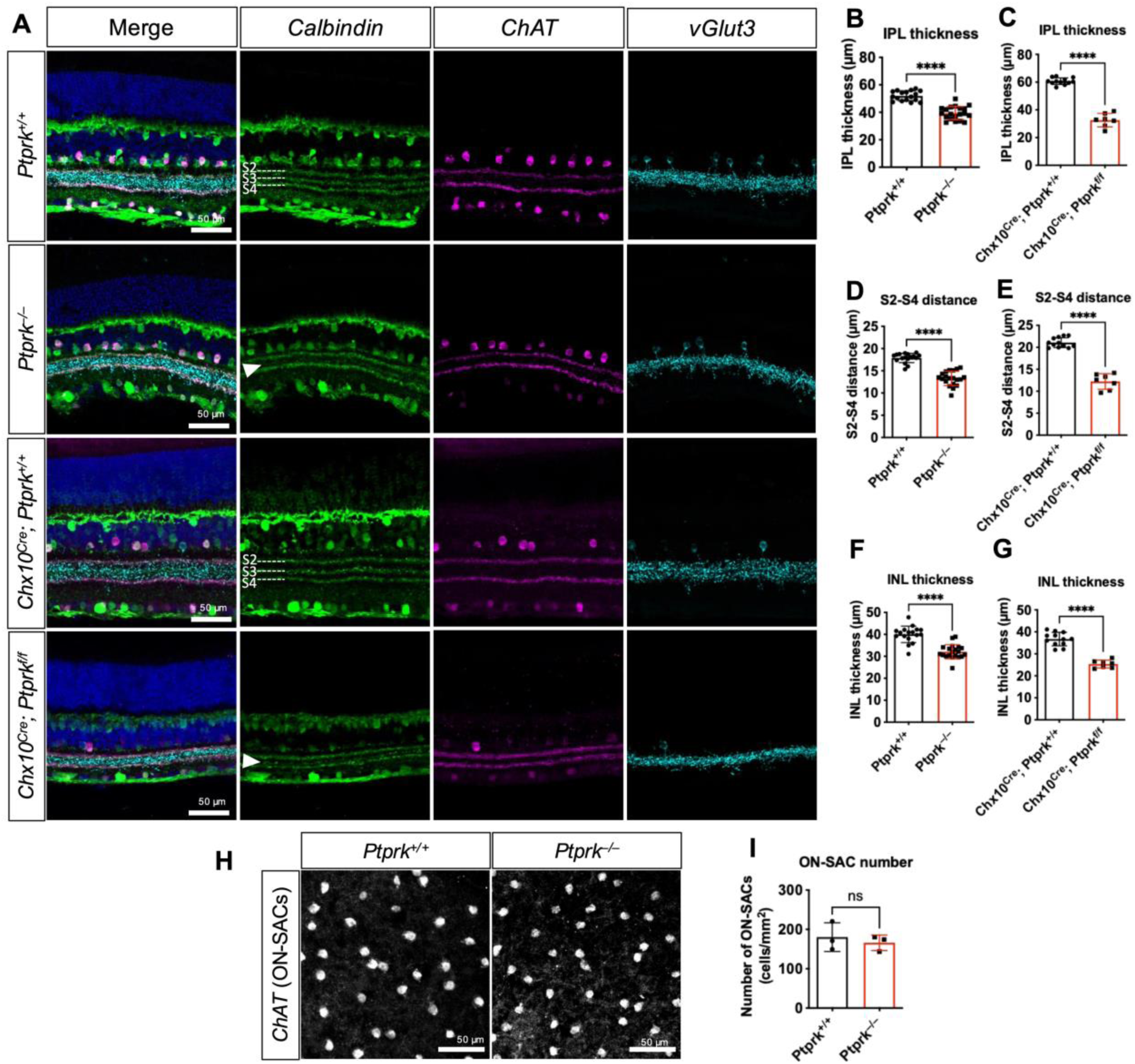
Retinal thinning in *Ptprk* mutants. **(A)** Cross-sections of adult retinas were immunostained with Calbindin (green), ChAT (magenta) and vGlut3 (cyan) to mark the retinal layers. Retinal stratification remained intact except for the missing Calbindin^+^ S3 in the mutants (indicated by white arrowheads). **(B-G)** Quantification of the thickness of IPL (B, C), S2-S4 (D, E) and INL (F, G) showed an overall thinning phenotype in both the global and pan-retinal knockouts. Sections chosen for quantification were as close to the center of the retina (i.e. close to the optic nerve) as possible while the lamial structures were still preserved. N = number of sections across 2-4 retinas. **(H-I)** count ON-SAC number on whole-mount adult retinas (*ChAT*^+^) showed no change in ON-SAC number between *Ptprk^−/−^* and the control. N = number of animals. Data are shown in mean ± SD. \*\*\*\**p* < 0.0001.

**Supple. Figure 4.**
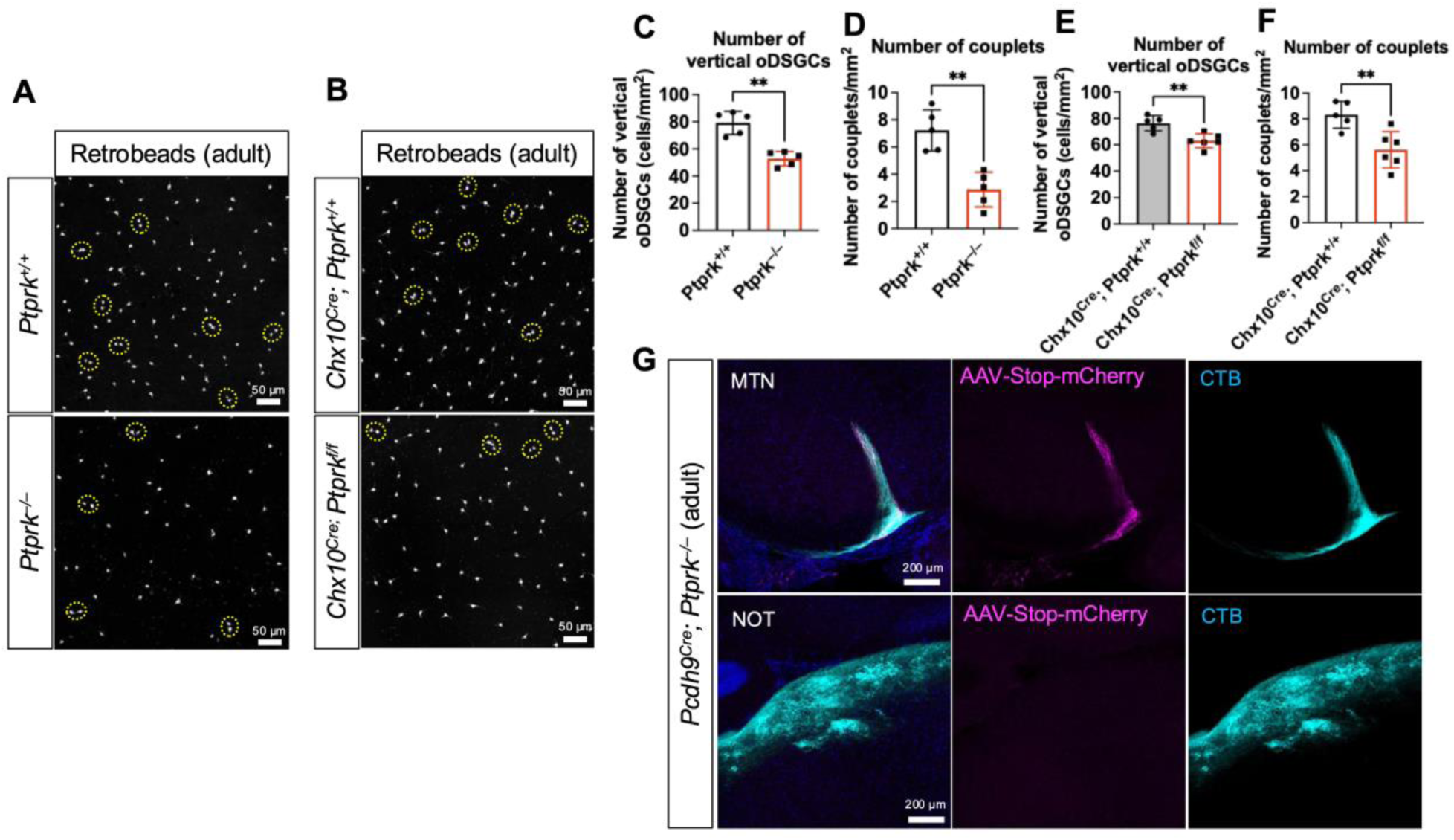
Quantification of couplets and Superior oDSGC central projection in *Ptprk* mutants. **(A, B)** Whole-mount adult retinas following retrograde injection of Retrobeads into the MTN of (A) global and (B) pan-retinal *Ptprk* mutants. Couplets, defined by the distance no more than 30 µm apart, are indicated by yellow dashed circles. **(C-F)** Quantification of the (C-D) global and (E-F) pan-retinal *Ptprk* mutants reveal fewer (C, E) Retrobead-labeled vertical oDSGCs as well as (D, F) couplets. N = number of animals. **(G)** Brain sections of MTN and NOT of adult *Pcdh9^Cre^; Ptprk^−/−^* mice that were intravitreally injected with CTB and the BChT virus (AAV-Stop-mCherry; used in Figure 3) to express mCherry in Superior oDSGC axons. While CTB delineated both MTN and NOT well; only MTN contained mCherry^+^ RGC axons.

**Supple. Figure 5.**
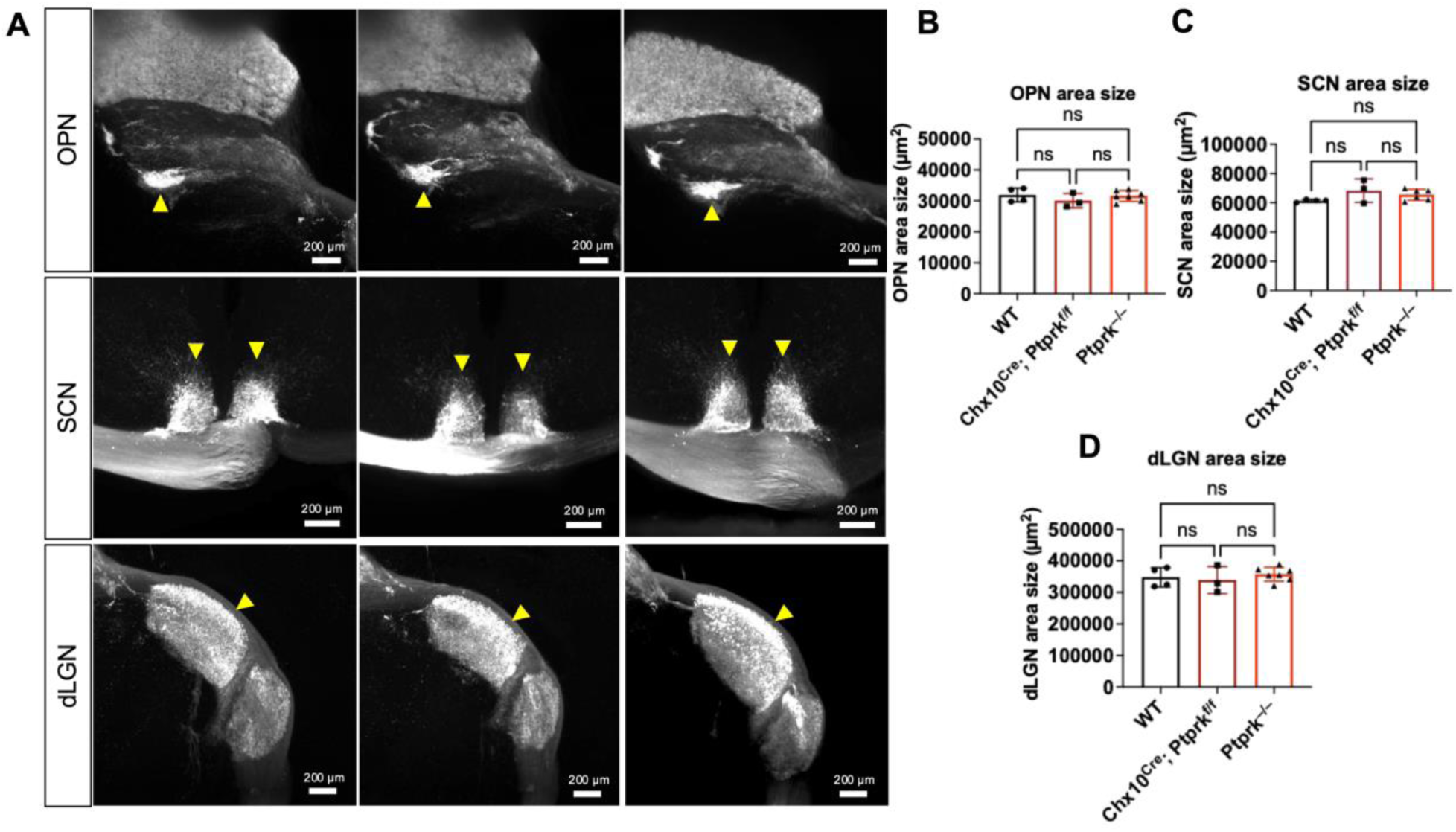
Innervation of other central nuclei. **(A)** Maximum intensity projections of light-sheet images of retino-recipient nuclei, labeled via intraocular injection of CTB. The olivary pretectal nucleus (OPN), suprachiasmatic nucleus (SCN) and dorsal lateral geniculate nucleus (dLGN) indicated by yellow arrowheads. **(B-D)** Quantification of the nuclei sizes from the projected 2D images of the (B) OPN, (C) SCN, and (D) dLGN.

**Supple. Figure 6.**
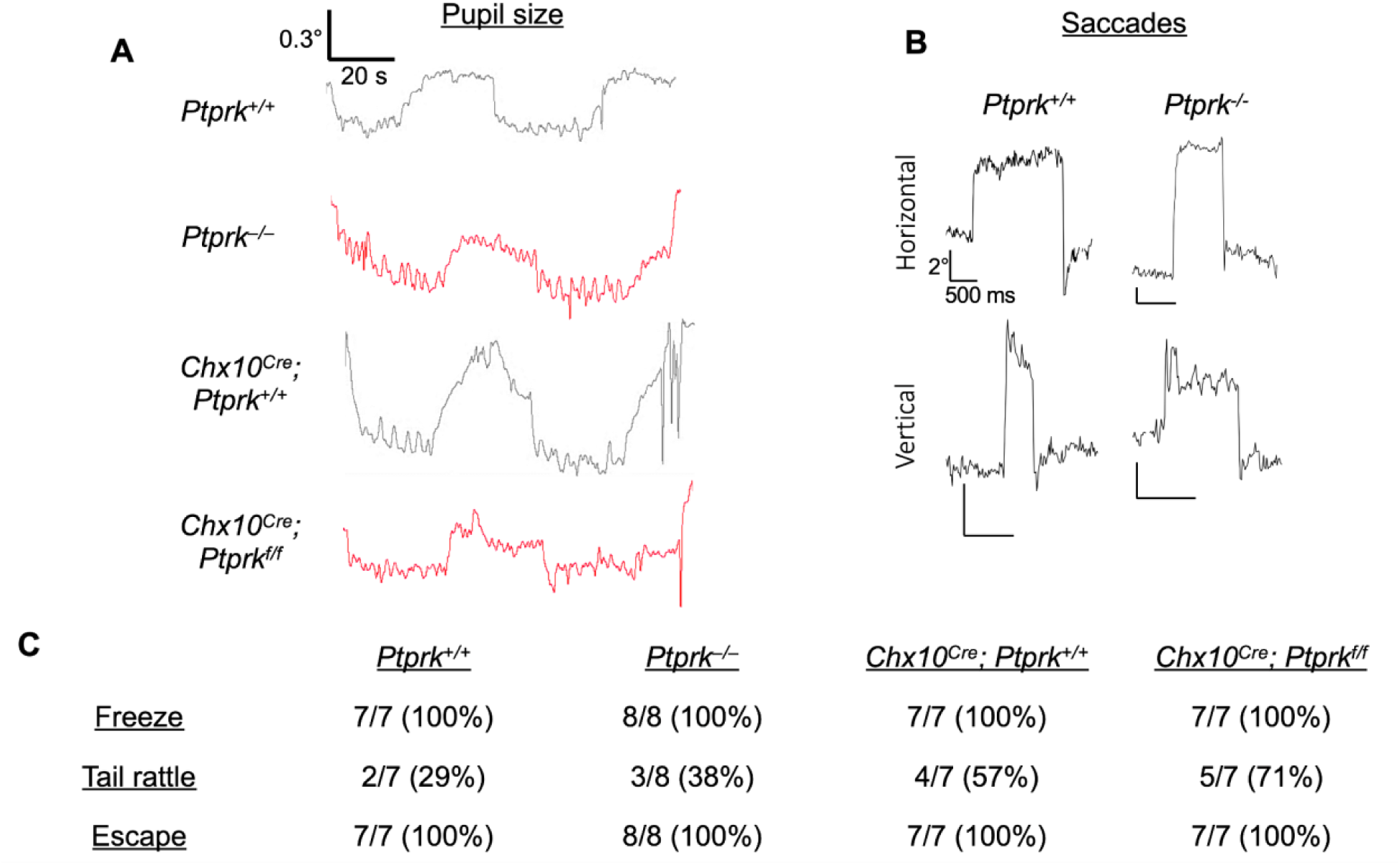
Other vision-related behaviors remain intact in *Ptprk* mutants. **(A)** *Ptprk^+/+^, Ptprk^−/−^, Chx10^Cre^; Ptprk^+/+^* and *Chx10^Cre^; Ptprk^f/f^* showed normal cycles of pupil dilation and constriction during OKR recordings, which consists of a 30-second light-on moving stimulus followed by a 30-second dark screen period. **(B)** Both WT and knockout animals showed normal voluntary horizontal and vertical saccades, defined by successful rapid eye movements in both horizontal (upper panels) and vertical (lower panels) axes. **(C)** Fear behaviors, including freezing, tail rattle, and escape, exhibited by the knockouts and their controls in response to overhead looming threats. All animals showed a normal fear response to overhead looming stimuli.

## Materials and Methods

All animal experiments were approved by the Institutional Animal Care and Use Committee (IACUC) at The Johns Hopkins University School of Medicine and at UCSF.

### scRNAseq data analyses

The analyses were conducted using the previously published datasets: P4-12 oDSGCs^15^, all P5 mouse RGCs^16^, adult mouse RGCs^1^, P19 mouse amacrine cells (ACs) ^41^, and human fetal RGCs^3^. *Ptprk* expression assessment using the P4-12 oDSGC dataset was performed as previously described^15^. The dot plots of gene expression of all adult RGCs and ACs were generated via the Broad Institute Single Cell Portal^1,40^. Assessment of gene expression using human fetal RGC dataset was done as previously described^3,19^.

### Head posting implantation

Adult mice that were at least 2 months of age were anesthetized using 2% isoflurane. Four1.00UNM × 0.120″ stainless steel screws were placed into the skull on both sides of the sagittal suture between the coronal and lambdoidal sutures. The screws were incorporated into a pedestal of dental cement (Ortho-Jet, Lang Dental; Wheeling, Illinois, USA) and an acrylic headpost—two M1.4 hex nuts embedded in dental cement (made in-house)—was placed on top of the pedestal. Mice were then tested for OKR response around 7 days after the surgery.

### OKR recording and analysis

Headposted mice were head-fixed in an animal holder (made in-house) inside a light-tight box. Four computer monitors were placed on each side of the box, displaying a black-and-white checkerboard patterns (stripe width 5°of visual angle). The animal’s anterior-posterior axis was aligned between two different pairs of monitors such that their two eyes saw visual rotational stimuli that emulated a virtual environment. A fixed infrared video camera inside the box recorded eye movements. Recordings of eye movements and presentation of checkerboard stimuli were custom-built in Matlab and LabView as previously described^15^. Two types of visual stimuli were used to elicit the OKR: sinusoidal stimuli with 0.2 Hz temporal frequency (5° of visual angle for 10 cycles per trial) and continuously rotating stimuli in 5°/s (30 s of rotating checkerboard followed by 30 seconds of grey screen; 10 cycles per trial). Stimuli were presented in both vertical and horizontal directions. Data were analyzed using Igor Pro (WaveMetrics, Portland, OR) as described previously^15^.

### Intravitreal CTB and AAV injections

Mice were anesthetized with 2% isoflurane. A 30G needle was used to create a hole at the corneal limbus. After the aqueous humor was gently squeezed out, 1 µl of CTB or AAV was injected through the hole with a mouth pipette. 0.75 µl of CTB and 0.75 µl of AAV were used for coinjections. The following viruses were used: AAV9 EF1a-BbChT (>1E+13; Addgene #45186-AAV9) and AAV2 CAG-mWGA-mCherry (2E+12; Virovek, Inc. Cat #117BUCSF89). The following CTB were used: CTB-488 (Thermo Fisher Cat # C34775), CTB-555 (Thermo Fisher Cat # C34778) and CTB-647 (Thermo Fisher Cat # C34778).

### Brain clearing and light sheet imaging

CTB-injected animals were transcardially perfused with phosphate buffered saline (PBS) with 0.5% heparin followed by perfusion with 4% paraformaldehyde (PFA) in PBS. Extracted brains were fixed in 4% PFA in PBS overnight at 4°C and later washed 3 times with PBS for 30 minutes. Brains were then dehydrated in 25%, 50%, 75%, and 100% methanol (MeOH) in H_2_O at room temperature for 30 minutes for each dilution. They were then washed again in 100% MeOH for 30 minutes, and then washed in a 66% dichloromethane (DCM)/33% MeOH solution for 3 hours at RT. The brains were then washed in 100% DCM at RT for 2 hr each until they sunk. They were then placed into dibenzyl ether (DBE) overnight. The following morning, DBE was replaced with fresh DBE and left to sit for at least 8 hours until the brains had significantly cleared. DBE was then replaced to ethyl cinnamate (ECi) overnight for imaging. Cleared brains were imaged on a Zeiss Lightsheet 7 microscope with a 5x objective in ECi with a refractive index of 1.558.

### Stereotaxic MTN injections

Adult mice were anesthetized using 2% isoflurane and secured in a stereotaxic apparatus. Lumafluor-555 retrobeads diluted 1:4 with ddH_2_O were injected into the MTN using a Hamilton Neuros syringe (PN: 65460-02) that was controlled by an automatic microsyringe controller (World Precision Instruments). The MTN was targeted using the following coordinates relative to bregma: anterior/posterior 0 mm, medial/lateral ± 0.85 mm, dorsal/ventral –5.4mm and –5.3mm with a needle tilt of 30° anteriorly. The needle was inserted to dorsal/ventral depth –5.5mm, raised 0.1mm for an injection of 500 nl in 10 nl/s, and raised 0.1mm again for another 500 µl injection. Following each injection, the needle stayed in place for 5 minutes to allow for diffusion of the retrobeads.

### Immunohistochemistry

Tissue, either eyes or brains, were extracted by transcardially perfusing the mice with PBS followed by 4% PFA and fixed in 4% PFA at 4°C overnight. For staining whole-mount retinas, the retinas were dissected and incubated for 2 days at 4°C with primary antibodies in PBS with 0.4% Triton X-100, 5% donkey (or goat, depending on the secondary antibody) serum. Retinas were washed 4 times for 1 hour each at room temperature with 0.1% Triton X-100 in PBS before incubated with secondary antibodies in PBS with. 0.1% Triton X-100 and 5% donkey (or goat) serum at 4°C overnight. Retinas were then washed 4 times, 1 hour each, at room temperature with 0.1% Triton X-100 PBS and mounted for imaging.

For staining retinal cross sections, cryoprotection was done by incubating punctured eyes in PBS containing 30% (w/v) sucrose overnight at 4°C. Eyes were frozen in Neg-50 (Kalamazoo, MI) and sectioned via a cryostat in 20 µm thickness. Retinal sections were blocked in PBS containing 5% donkey (or goat) serum and 0.4% Triton X-100 at room temperature for 1 hour followed by incubation with primary antibodies in PBS containing 5% donkey (or goat) serum and 0.4% Triton X-100 overnight at 4°C. Sections were washed 6 times for 5 minutes each at room temperature with PBS containing 0.1% Triton X-100 followed by incubation of secondary antibodies in PBS containing 5% donkey (or goat) serum and 0.4% Triton X-100 at room temperature for 1 hour. Sections were washed 6 times for 5 minutes each at room temperature with PBS and 0.1% Triton X-100 and then mounted for imaging.

For staining brain cross sections, brains were embedded in PBS containing 4% (w/v) low-melting-point agarose and sliced into 150 µm sections using a vibratome. Sections were permeabilized in permeabilization solution (3% BSA, 0.3 Triton X-100 and 0.02 sodium azide in PBS) at room temperature for 30 minutes followed by incubation with primary antibodies in permeabilization solution with 5% goat serum at 4°C overnight. Sections were washed 3 times for 1 hour each at room temperature with PBS containing 0.1% Triton X-100 followed by incubation of secondary antibodies in PBS containing 0.1% Triton X-100 at 4°C overnight. After washing 3 times for 1 hour each with 0.1% Triton X-100 PBS, sections were mounted for imaging.

Primary antibodies used in this study include Rabbit anti-dsRed (Living Colors, 1:1000), Chick anti-GFP (AVES, 1:1000), Rabbit anti-Rbpms (Proteintech, 1:500), Guinea Pig anti-vGlut3 (Synaptic Systems, 1:1000), Rabbit anti-Calbindin (Swant, 1:1000), and Goat anti-ChAT (Abcam, 1:250). All images were taken with a Zeiss LSM 700 confocal microscope.

### Electrophysiology

Retrobeads were visualized in oDSGCs during electrophysiology recordings using two-photon imaging at least two days following injections of retrobeads into the MTN. Prior to electrophysiology, mice were dark-adapted overnight. Mice were euthanized by cervical dislocation and eyes were enucleated under infrared light. Because retrograde tracers were injected into one MTN, the contralateral eye was collected. Retina dissections were performed in the dark using infrared converters on a dissecting microscope and whole-mounted on a glass coverslip. Following enucleation, brains were harvested and fixed in 4% PFA, then sectioned using a vibratome to confirm the presence of the retrobeads in the MTN. Retinas were perfused with warmed (30-35 deg C), oxygenated (95% O_2_/5% CO_2_) bicarbonate-based Ames solution during the dissection and the recording. Prior to recording from ganglion cells, twelve locations around the perimeter of the retina, defined by cuts made along dorsal-ventral and nasal-temporal axes, were measured to calculate the geometric center of the tissue. The center was used to establish a coordinate system on the retina and measure distances between recorded ganglion cells.

Cell-attached recordings of oDSGCs were performed as described in (Harris and Dunn, 2023). Retrobead fluorescence in retinal ganglion cell somas were identified using a two-photon microscope (peak emission at 860 nm) and cells were recorded using patch electrodes (Sutter) filled with HEPES-buffered Ames using IR light. Labeled retinal ganglion cells located within 30 µm of each other were defined as couplets. Data was acquired at 10kHz with a MultiClamp 700B amplifier using Symphony 2 DAS, run through Matlab software. Visual stimuli were presented using a DLP projector (peak emission at 405 nm), centered on the cell somas and covered a 427 x 311 µm region. Responses to five repetitions of the moving bar stimuli (3.2° in width), moving at 10°/sec in 8 directions (every 45°), were collected for each cell.

All electrophysiology data was analyzed using Matlab software. Responses were averaged across five trials per stimulus direction, generating a representative tuning curve for each cell. Preferred directions for each cell were calculated as the direction of the vector sum of spike responses to all stimulus directions. Normalized tuning curve area was calculated as the tuning curve area divided by the response in the preferred direction. The direction selectivity index (DSI) was calculated as the vector sum of responses divided by the scalar sum of responses to all stimulus directions.

